# Consensus Formation History Shapes Collective Updating

**DOI:** 10.64898/2026.09.23.753723

**Authors:** Chuyang Sun, Ruoxi Zhou, Dandan Zhang, Jialu Wang, Yu Meng, Rixin Tang

**Author notes:** **Corresponding Author:** Rixin Tang, PhD. Nanjing University. These authors contributed equally to this work.

## Abstract

Consensus is rarely a one-off achievement. In a changing world, groups must repeatedly update judgments they have already agreed upon, making consensus updating a common feature of real-world collective decisions. However, previous research on collective decision-making has focused mainly on how groups reach a single consensus, leaving the updating of established consensus largely unexamined. This raises two unanswered questions: does consensus updating involve biases analogous to sunk cost, and are such biases related to the cost of the previous consensus? Building a Fake Consensus paradigm, we manipulated consensus-formation cost and examined how dyads updated a subjectively established consensus after new information arrived. An independent individual experiment provided information-matched updating baselines, allowing us to determine whether dyads updated less or more than individuals responding to the same evidence. Contrary to sunk-cost predictions, low-cost consensus led dyads to update less than the individual baseline, whereas medium- and high-cost consensus led them to update more. The pathway through perceived consensus sharedness was significant, and the association between sharedness and updating was stronger at higher left temporoparietal junction inter-brain synchrony. These findings indicate that the history of consensus formation shapes subsequent updating by altering the psychological meaning of shared endorsement, while interpersonal alignment determines how this shared experience translates into coordinated behavior. To our knowledge, this is the first experimentally controlled study of consensus updating in teams to isolate formation history from actual disagreement, providing a new paradigm for studying how collective judgments change after agreement.

## 1. Introduction

Reaching consensus about where to go and when to act is a precondition for coordinated action and, in many species, for survival (Krause & Ruxton, 2002; Conradt & Roper, 2005). Yet consensus is rarely a one-off achievement. The world keeps changing, and groups must repeatedly revise judgments they have already agreed upon. Consider a board that has painfully settled on a strategy: when the market turns and new data arrive, does it update, or does it hold on to a hard-won position? Research on collective decisions has illuminated how groups reach agreement (Suzuki et al., 2015; Kerr & Tindale, 2004; Bahrami et al., 2010; Deng et al., 2026). A further question concerns the psychological legacy of that process: does working harder to reach consensus make a judgment more resistant to updating, or can the same difficult history weaken its hold? Answering this question requires explaining how the experience of reaching agreement changes the meaning of that agreement for subsequent decisions.

Consensus formation supplies two potentially competing movement directions for the earlier judgment. First, it requires an investment of time and effort. Sunk-cost and escalation-of-commitment research suggests that prior investments can encourage persistence with an existing course of action, including in groups (Arkes & Blumer, 1985; Whyte, 1993; Staw, 1976; Sleesman et al., 2012). A straightforward investment account therefore predicts less updating after a more costly consensus. On the contrary, consensus formation provides information about others’ preferences and their insistence on prior choices (Suzuki et al., 2015). Such social information matters because a sense of shared understanding can provide an epistemic basis for confidence in a judgment (Echterhoff & Higgins, 2017; Higgins et al., 2021; Echterhoff et al., 2009). Building on these findings, we propose that repeated failures to agree may weaken perceived joint endorsement, even when agreement is eventually declared. A costly agreement may therefore provide less reassurance and remain more open to updating. This account raises the possibility that the social meaning acquired during negotiation can counteract the stabilizing influence of prior investment.

We propose that perceived consensus sharedness provides one route from formation history to subsequent updating. We define sharedness as the extent to which members feel that the established judgment represents both of them. This proposal draws on shared-reality research (Rossignac-Milon et al., 2021), which links an experienced common understanding to the social validation of judgments and subjective certainty (Echterhoff & Higgins, 2017; Higgins et al., 2021). Related work shows that confidence can favor the processing of evidence consistent with an existing decision (Rollwage et al., 2020; Kappes et al., 2020). Applied to consensus updating, easily reached agreement may leave members with a stronger sense that their judgment is jointly supported, whereas repeated disagreement may make the eventual outcome feel provisional. We therefore expect greater formation cost to reduce perceived sharedness and lower sharedness to be associated with greater subsequent updating. Sharedness is the measured construct in this account; shared reality and epistemic certainty provide theoretical reasons for expecting it to matter.

We further propose that the behavioral relevance of sharedness depends on interpersonal alignment developed during consensus formation. Feeling that a judgment represents both members and coordinating one’s response with a partner are distinguishable aspects of a joint decision. Alignment may help members continue to rely on a position they experience as shared, while also helping them make compatible updatings when new evidence arrives. Work on brain-to-brain coupling motivates inter-brain synchrony (IBS) as a candidate neural measure of coordinated processing during interaction (Hasson et al., 2012; Redcay & Schilbach, 2019; Czeszumski et al., 2022; Reinero et al., 2021). We focused on the left temporoparietal junction (lTPJ), informed by evidence implicating temporoparietal regions in reasoning about others’ perspectives (Schurz et al., 2014; Saxe & Kanwisher, 2003; Samson et al., 2004; Jiang et al., 2015). Our prediction concerns moderation: the negative association between sharedness and update magnitude should be stronger at higher IBS. A complementary behavioral test asks whether IBS predicts more similar updates between partners, especially when substantial information change requires substantial updating. Neither prediction requires formation cost to change lTPJ IBS itself.

To test these, we introduce a new paradigm called ‘Fake Consensus’ paradigm (for details, please find Method section). In this paradigm, there are two stages of consensus: stage 1 manipulated the number of consensuses rounds to control the consensus cost, and set the last round of individuals’ choices as their own perceived team consensus (fake consensus). After rating how strongly this judgment represented both members, participants received new information and attempted to agree on how far to revise it, but only the first post-change response was analysed. Their first responses, made before renewed agreement feedback, indexed initial updating; simultaneous fNIRS recordings during formation provided IBS estimates. This design separated assigned formation history from naturally occurring disagreement and distinguished three outcomes: updating relative to an empirical individual baseline, perceived sharedness, and the discrepancy between members’ signed updates.

By using this paradigm, we found that formation history changed the direction of dyads’ deviation from the individual baseline: dyads updated less after low-cost consensus and more after medium- and high-cost consensus. Lower perceived sharedness statistically accounted for part of the association between greater cost and greater updating. The estimated sharedness–updating association was also stronger at higher lTPJ IBS, although the interaction’s statistical support depended on the inference method. Higher lTPJ IBS predicted more similar initial updatings when information changed substantially, providing complementary evidence for its relevance to coordinated updating. Together, these findings support an account in which formation history shapes how jointly held a judgment feels, and alignment helps determine how that subjective experience relates to later behavior.

## 2. Experiments

### 2.1 Experiment 1: Individual updating baselines

The study comprised two experiments. Experiment 1 established information-specific individual updating baselines (Bahrami et al., 2010; Deng et al., 2026). Experiment 2 used an fNIRS hyperscanning design (Montague et al., 2002) to examine how experimentally assigned consensus-formation cost was associated with subsequent dyadic updating. All participants provided written informed consent. The protocol was approved by the Psychology Research Ethics Committee at Nanjing University (NJUPSY2025007002). Because Experiment 2 used false performance feedback, participants were debriefed after the session.

#### 2.1.1 Participants

Seventy-seven adults participated in Experiment 1 (age: *M* = 24.04 years, *SD* = 2.72; 41 women). None participated in Experiment 2.

#### 2.1.2 Task and procedure

Participants completed an individual rainfall-forecasting task. On each trial pair, they viewed temperature, humidity, and cloud-cover information and rated expected rainfall on a 7-point scale (1 = no rain, 7 = extremely heavy rain). Updated information then indicated that humidity would increase or decrease by a small, moderate, or large amount, after which participants made a second forecast. The experiment comprised 18 trial pairs, with six at each information-change level. Increase and decrease directions were balanced within level, and trial order was randomized. Each pair consisted of 30 s of initial-information viewing, a 7-s forecast, 30 s of updated-information viewing, and a second 7-s forecast (Figure 1).

**Figure 1.**
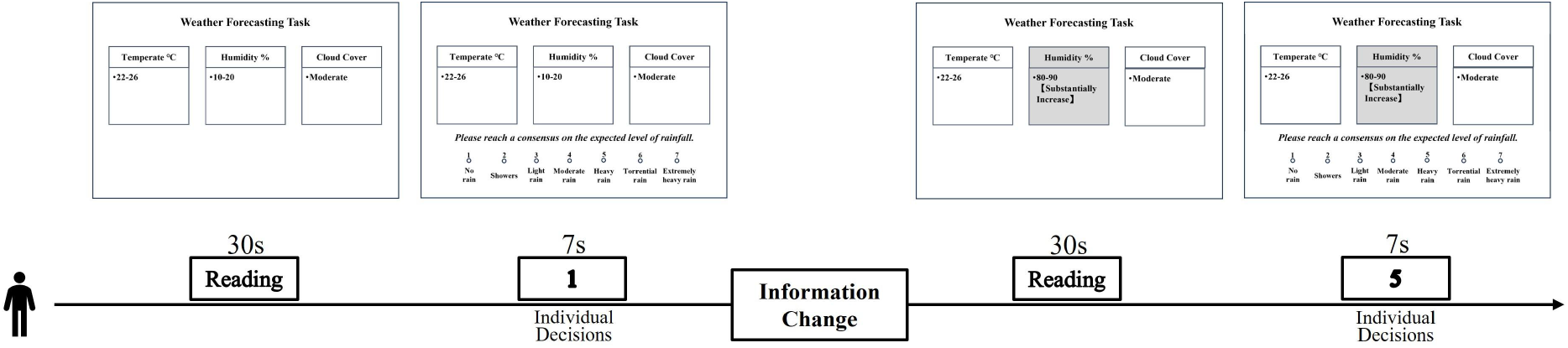
Experimental procedure for establishing individual updating baselines. Participants independently judged expected rainfall from meteorological information and revised their forecasts following a low, medium, or high change in humidity. Mean absolute changes in judgment at each information-change level provided the corresponding individual updating baselines for Experiment 2. The diagram illustrates a single trial pair. Shading identifies the updated information, and numerical responses are illustrative.

#### 2.1.3 Analysis

The primary measure was the absolute difference between the two forecasts. Mean update magnitude was calculated for each participant at each information-change level and analyzed with a one-factor repeated-measures analysis of variance. Mauchly’s test assessed sphericity, and planned paired comparisons quantified separation among levels. The three means served as the empirical individual updating baselines (*I*⁎) for Experiment 2. These values describe typical updating in the same task without prior dyadic consensus.

#### 2.1.4 Results

Information-change magnitude affected individual updating, *F*(2, 152) = 47.58, *p* < .001, generalized η² = .166. Sphericity was not violated, Mauchly’s *W* = .971, *p* = .334. Mean update magnitude increased from low (*M* = 0.675, *SD* = 0.475) to medium (*M* = 1.123, *SD* = 0.687) to high information change (*M* = 1.346, *SD* = 0.701; Figure 2). All planned comparisons were significant: low versus medium, *t*(76) = 6.57, *p* < .001, *d* = 0.75; medium versus high, *t*(76) = 3.38, *p* = .001, *d* = 0.39; and low versus high, *t*(76) = 8.87, *p* < .001, *d* = 1.01. Direction-coded updates showed the same ordering (0.52, 1.02, and 1.22). Accordingly, 0.675, 1.123, and 1.346 were used as the low-, medium-, and high-information baselines in Experiment 2.

**Figure 2.**
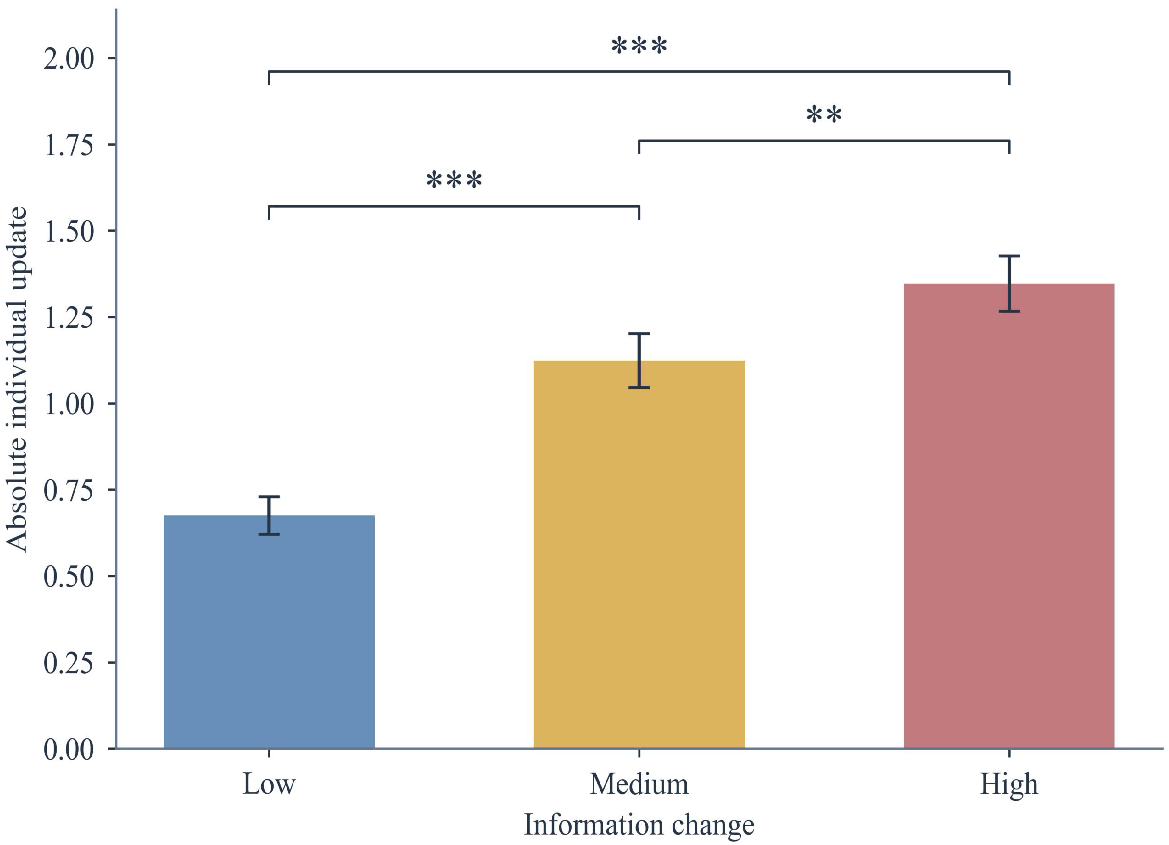
Individual updating baselines in Experiment 1. Mean absolute changes between the two forecasts are shown for each information-change level (*N* = 77). Error bars indicate SE. Brackets indicate planned paired comparisons: ** *p* < .01; *** *p* < .001.

### 2.2 Experiment 2: Consensus formation and dyadic updating

#### 2.2.1 Participants and design

Forty-two adults were tested in 21 dyads (age: *M* = 22.44 years, *SD* = 3.00; 34 women). Dyad members were strangers and were randomly assigned to positions A and B. All 21 dyads were included in the behavioral and IBS analyses. The experiment used a 3 × 3 within-dyad design crossing consensus-formation cost (low, medium, high) with information change (low, medium, high). Each dyad completed 18 blocks, with six blocks at each marginal level of cost and information change. Meteorological parameters were sampled with replacement from 12 predefined sets to preserve trial-to-trial variation within a controlled stimulus pool.

#### 2.2.2 Consensus formation and updating task

The task was designed to test how assigned consensus-formation history influenced the updating of a previously accepted judgment. Dyads were instructed to form a reasonable consensus as quickly as possible and were told that bonus payment would depend on their actual responses. Each block comprised two stages (Figure 3). In Stage 1, participants viewed the initial meteorological information for 30 s; this information remained visible while they responded. On each round, participants selected one of seven response buttons during a 7-s window and then received 1.5 s of manipulated feedback about whether the dyad had reached consensus.

**Figure 3.**
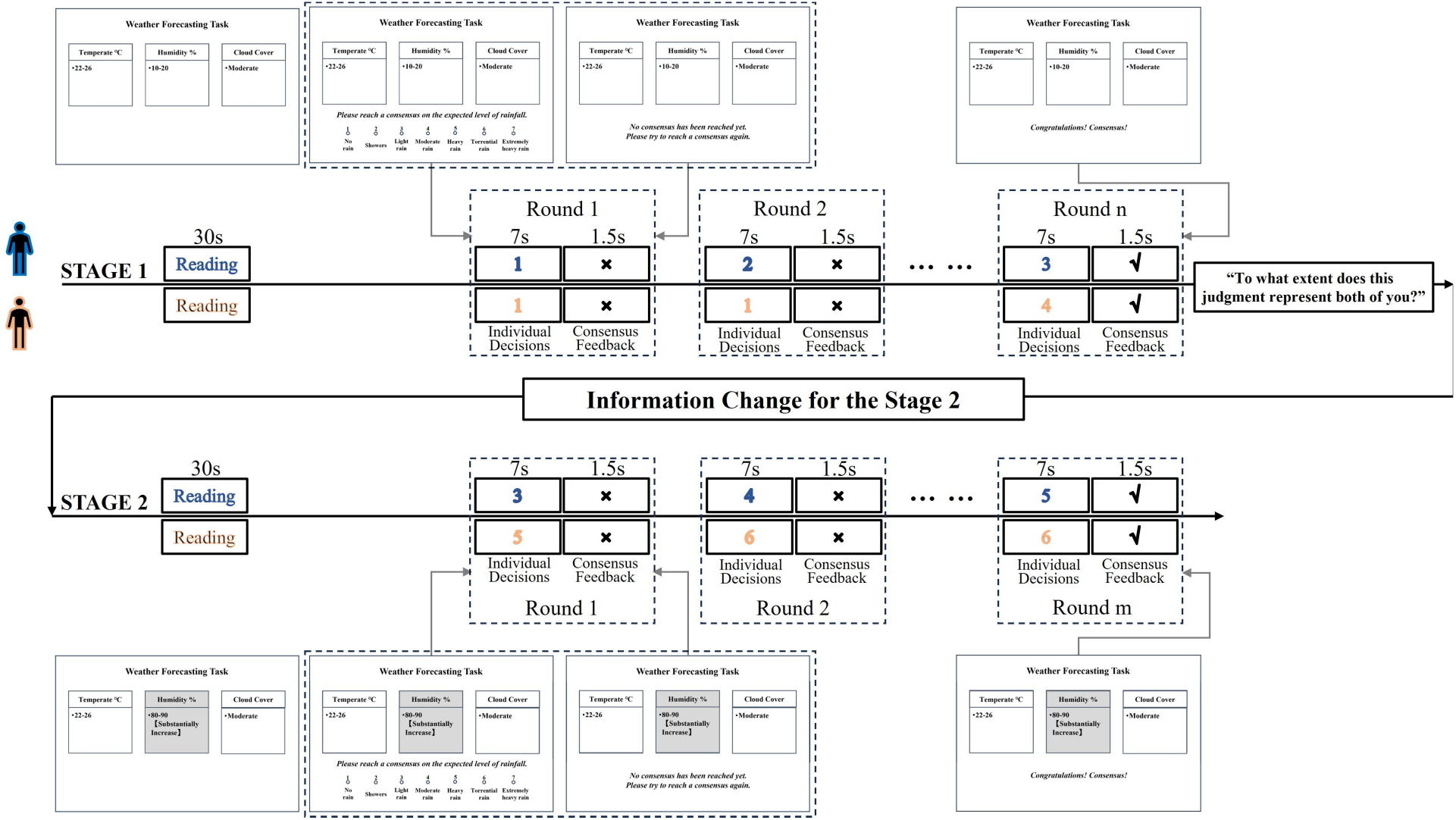
Two-stage consensus formation and updating task in Experiment 2. Blue and orange distinguish the two members; crosses and check marks indicate feedback signaling disagreement and consensus, respectively. The symbols n and m denote the final rounds of Stages 1 and 2. Numerical responses are illustrative.

The program predetermined the round on which consensus would be declared: rounds 1–3 in the low-cost condition, 4–6 in the medium-cost condition, and 7–9 in the high-cost condition, with the target round sampled within the relevant interval. Before the target round, both members were told that their predictions were inconsistent regardless of their responses. On the target round, both were told that consensus had been reached. The program separately stored each member’s final Stage 1 forecast as that member’s perceived consensus. The two stored values could differ; the program did not require both members to occupy the same response-scale position. Thus, assigned cost combined the number of response rounds, elapsed negotiation time, and repeated disagreement feedback; these components were not manipulated separately.

Immediately after consensus was declared and before new information appeared, each participant rated perceived consensus sharedness by answering “To what extent does this judgment represent both of you?” on a 1–7 slider (1 = not at all representative; 7 = completely representative) within 2 s. The mean of the two ratings indexed dyadic sharedness. This item measured the perceived joint representativeness of the judgment; it did not separately assess confidence in its accuracy, epistemic certainty, or a broader experience of shared reality.

In Stage 2, the changed meteorological parameter—for example, a change in humidity—was visually highlighted and remained on screen for 30 s. Participants then completed further 7-s response rounds followed by 1.5-s feedback. Renewed consensus required equal signed changes from each member’s own perceived consensus: moves from 2 to 4 and from 5 to 7 both counted as +2 and met the criterion, whereas moves from 1 to 2 and from 7 to 6 did not. Agreement therefore concerned the direction and magnitude of updating, even when final numerical judgments differed. The primary analyses used the first Stage 2 decisions, made after the information changed but before participants received further agreement feedback. These decisions captured initial updating within a continuing joint task and could already incorporate expectations about the partner.

#### 2.2.3 Data acquisition

Behavioral and neural data were recorded simultaneously throughout Experiment 2. The two members of each dyad completed the task through separate response interfaces. The experimental program independently recorded each member’s judgment together with the corresponding dyad, block, experimental stage, interaction round, consensus-cost condition, and information-change condition. The onset and offset of the principal task events—including information presentation, judgment, feedback, declared consensus, and new-information presentation—were also recorded. These event timings were used to align the behavioral records with the continuous fNIRS signals and to define the Stage 1 consensus-formation windows for subsequent neural analysis.

Cortical haemodynamic activity was recorded simultaneously from both dyad members using a LABNIRS continuous-wave functional near-infrared spectroscopy system (Shimadzu Corporation, Kyoto, Japan). The system used three wavelengths of near-infrared light (780, 805, and 830 nm) and acquired signals at a sampling rate of 30.3 Hz. Changes in oxygenated haemoglobin (HbO), deoxygenated haemoglobin (HbR), and total haemoglobin (HbT) concentrations were derived according to the modified Beer–Lambert law (Cope & Delpy, 1988). Subsequent neural analyses focused on HbO signals.

Each participant wore a custom 25-channel montage positioned according to the international 10/05 system (Jurcak et al., 2007), with a source–detector separation of 30 mm. Corresponding probe arrangements were used for the two participants, enabling IBS to be calculated between homologous channels. The montage covered bilateral dorsolateral prefrontal and temporoparietal regions (Czeszumski et al., 2022) and was divided into five regions of interest: the left temporoparietal junction (lTPJ), right temporoparietal junction (rTPJ), anterior dorsal prefrontal cortex (aDPFC), mid-dorsal prefrontal cortex (mDPFC), and posterior dorsal prefrontal cortex (pDPFC) (Badre & D’Esposito, 2009). The lTPJ was the literature-informed primary ROI (Jiang et al., 2015; Samson et al., 2004), and the other regions provided comparison measures (Cui et al., 2012; Czeszumski et al., 2022; Tang et al., 2016). The probe arrangement and channel locations are shown in Figure 4.

**Figure 4.**
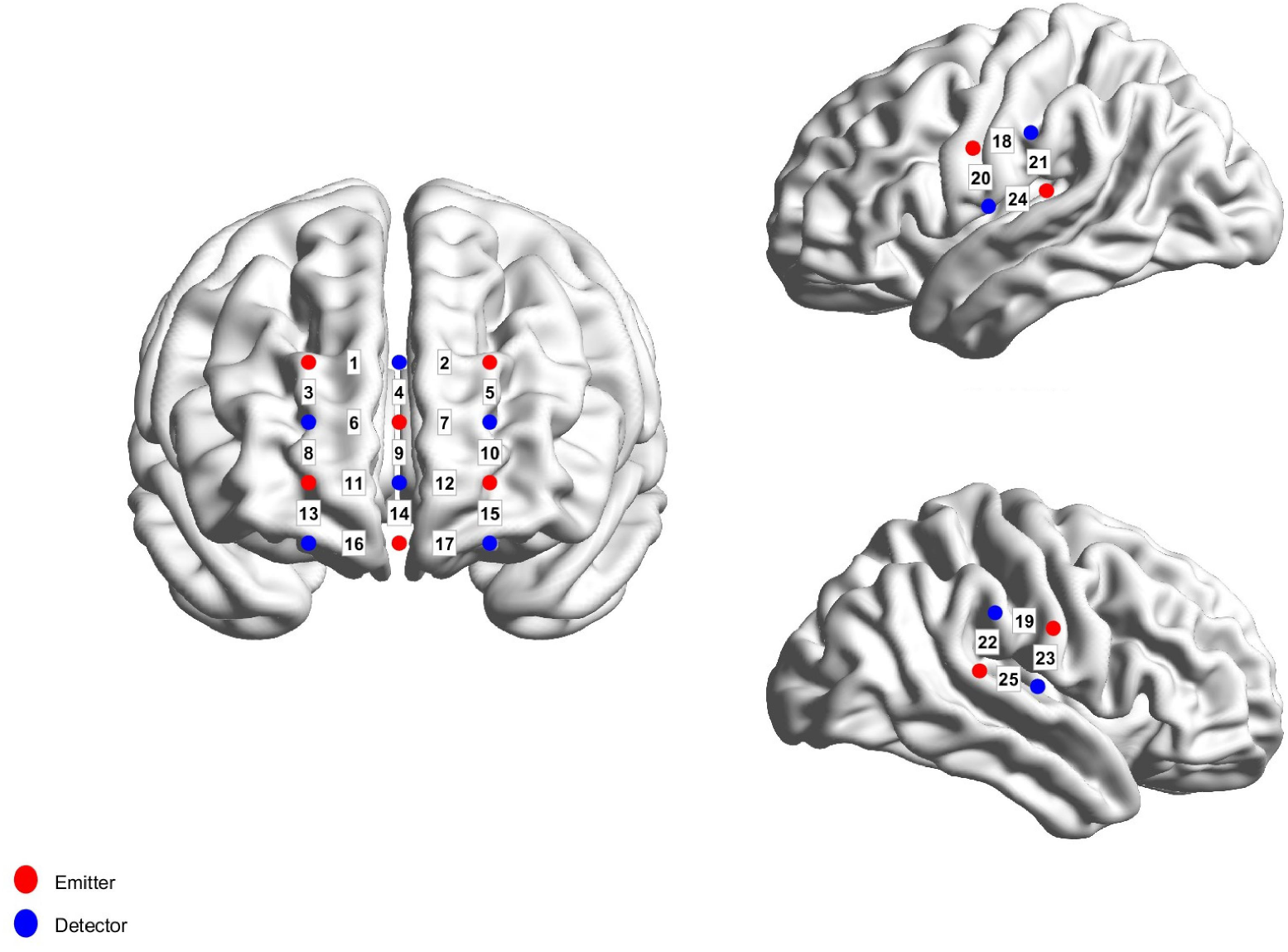
Optode arrangement and channel locations for fNIRS hyperscanning. The functional near-infrared spectroscopy (fNIRS) montage comprised 25 measurement channels per participant, covering bilateral dorsolateral prefrontal and temporoparietal regions. Red and blue circles indicate light sources and detectors, respectively, and numbers identify measurement channels.

The spatial positions of the channels were measured using a Polhemus FASTRAK three-dimensional digitizer. Channel locations were registered to Montreal Neurological Institute space using the virtual spatial registration procedure implemented in the NIRS-SPM toolbox, with Nz, Cz, AL, and AR as anatomical reference points (Singh et al., 2005; Tsuzuki et al., 2007). The MNI coordinates, corresponding Brodmann areas, and ROI membership of individual channels are provided in **Table S1**.

#### 2.2.4 Data Analysis

##### 2.2.4.1 Behavioral data analysis

###### Behavioral outcomes

For member i, the signed first-round movement was Δᵢ = first Stage 2 decision − perceived consensus. The primary outcome, baseline-adjusted updating (change), was the dyad’s mean absolute movement minus the information-matched individual baseline: change = (|Δ*_A_*| + |Δ*_B_*|)/2 − I*. Here, I* was the Experiment 1 mean absolute update for the corresponding information-change level. Positive and negative values indicated more and less updating than the individual baseline, respectively. Absolute movement indexed the amount of updating irrespective of direction on the response scale; it did not measure updating accuracy.

The between-member updating discrepancy, gap, was |Δ*_A_* − Δ*_B_*|. Smaller values indicated more similar signed changes from the members’ respective perceived consensuses. A gap of zero therefore indicated agreement about how to revise, without requiring identical final judgments.

The analysis unit was the dyad × cost × information cell. Measures were first computed for each block and then averaged across blocks within a cell; repeated blocks from the same cell were not treated as independent observations. The analyses first tested the effect of formation cost and the sharedness pathway, then examined IBS as a correlate of coordinated updating and a moderator of the sharedness–updating association.

###### Cost effects and sharedness pathway

Categorical mixed-effects models predicted each behavioral outcome from consensus cost, information change, and their interaction, with a dyad random intercept. Dyad-level means averaged across available information conditions were compared with zero to locate updating relative to the individual baseline; cell-specific comparisons used Holm adjustment. Auxiliary models examined starting judgments, available response-scale range, Stage 1 round count, task progression, and partners’ earlier adjustments. For gap, mean absolute movement and directional headroom were added to assess whether any cost association extended beyond movement magnitude and scale constraints.

We then tested whether perceived consensus sharedness accounted for part of the association between consensus cost and subsequent updating. The path models were estimated using ordinary least squares, with an intercept and linear scores for consensus cost and information change coded −1, 0, and 1. The cost-to-sharedness path (a) was estimated by regressing sharedness on cost and information change. The total cost association (c) was estimated by regressing change on cost and information change; adding sharedness provided the sharedness-to-updating path (b) and the direct cost association (c′). Single-coefficient inference used dyad-clustered CR2 covariance estimates with coefficient-specific Satterthwaite degrees of freedom (Bell & McCaffrey, 2002; Pustejovsky & Tipton, 2018). The indirect association was quantified as ab and evaluated using a joint-significance test of the two component paths, with *p*_joint_ = max(*p_a_*, *p_b_*) (MacKinnon et al., 2002; Yzerbyt et al., 2018). This test rejected the null hypothesis of no indirect association only when both paths were significant. A confidence interval based on 5,000 dyad-level bootstrap samples was retained as a sensitivity analysis. These path estimates describe an indirect association; sharedness was measured rather than experimentally manipulated (Spencer et al., 2005).

To examine whether the cost-related updating pattern was present in both members, we conducted an auxiliary member-level analysis using direction-aligned calibration errors, *eᵢ* = dir × Δᵢ − *I*⁎, where dir was +1 or −1 according to the direction indicated by the new information. Within each block, the member with the smaller |*eᵢ*| was assigned to the near role and the member with the larger |*eᵢ*| to the far role. These roles were assigned separately within each block and did not represent fixed participant identities. The resulting errors, *e*_near_ and *e*_far_, were averaged within each dyad × cost × information cell, and their linear associations with consensus cost were estimated separately. Because sorting members by absolute error can influence the estimated slopes, the observed slopes were compared with an unsorted reference distribution generated from 1,000 random assignments of members A and B to the two member series within each block, followed by cell-level aggregation and slope estimation.

##### 2.2.4.2 Neural data analysis

Neural analyses were conducted on the preprocessed HbO signals using MATLAB. First, the initial trial (practice trial) was removed from each session, followed by retention of only correct response trials, excluding error trials. Subsequently, a 3-standard deviation criterion was applied to exclude data points beyond the mean ± 3 standard deviations range to ensure data quality. The time series were resampled to 1 Hz, and only the Stage 1 consensus-formation period was used to calculate inter-brain synchrony.

Wavelet transform coherence (WTC) was computed between homologous channels of the two dyad members using a Morlet wavelet (Grinsted et al., 2004; Cui et al., 2012). For the HbO time series *x(t)* and *y(t)*, squared wavelet coherence was calculated as

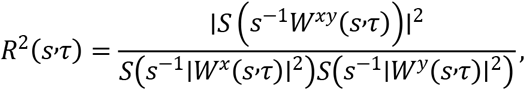

where *W^x^* and *W^y^* denote the continuous wavelet transforms of the two signals, *W^xy^* =*W^x^W^y^*\* denotes their cross-wavelet spectrum, *s* denotes wavelet scale, *τ* denotes time, and *S* denotes smoothing across time and scale.

For each dyad, homologous channel pair, consensus-cost condition, and frequency bin, *R*^2^ values were first averaged across the Stage 1 time points belonging to that condition. The resulting time-averaged values were converted to coherence and Fisher-*z*-transformed. The Fisher-*z*-transformed spectra were then averaged across all homologous channels to obtain a whole-channel mean spectrum for each dyad and cost condition. To identify task-related frequency ranges, a one-way repeated-measures ANOVA was conducted separately at each frequency bin, with consensus-formation cost (low, medium, and high) as the within-dyad factor. This frequency-screening analysis tested only the main effect of consensus cost.

Candidate frequency clusters were required to contain at least three adjacent frequency bins showing significant main effect (p < .05) on consensus cost. The statistic for each cluster was the sum of the F values across its constituent bins. Cluster-level significance was evaluated using 1,000 permutations in which the three consensus-cost labels were randomly reassigned within each dyad. For each permutation, the largest cluster statistic was retained to form the null distribution, and the cluster-level p value was calculated as the proportion of permuted maximum statistics equal to or greater than the observed statistic. Frequency clusters overlapping systemic physiological ranges were not carried forward (Tachtsidis & Scholkmann, 2016). The selected frequency range and cluster statistic are reported in the Results.

The same selected frequency band was applied to all ROIs to obtain their Stage 1 IBS estimates. The lTPJ was the literature-informed primary ROI; rTPJ, aDPFC, mDPFC, pDPFC, and the whole-channel mean were comparison measures. Selection used the whole-channel cost effect, without using the subsequent behavioral outcomes or their interactions with IBS. Consequently, the within-band cost effect is descriptive of a band selected on that contrast and is not an independent confirmation of a neural cost effect (Kriegeskorte et al., 2009).

##### 2.2.4.3 Neural-behavioral modeling

IBS was centered separately within each consensus-cost condition to express neural variation relative to that condition’s mean. We first tested its behavioral relevance to coordinated updating: gap was regressed on consensus cost, information change, centered IBS, and the information-change × IBS interaction, controlling for mean absolute movement and directional headroom. This model tested whether IBS was associated with more similar signed updates when information changed, beyond differences in overall movement and the available scale range.

We next tested whether IBS moderated the association between perceived sharedness and baseline-adjusted updating. The model predicted change from consensus cost, information change, perceived sharedness, centered lTPJ IBS, and the sharedness × lTPJ IBS interaction. Conditional sharedness slopes were estimated at the mean of centered IBS and at one standard deviation below and above it. The indirect-association and moderation models were fitted separately. Their joint interpretation distinguishes a subjective pathway from a neural moderator; it does not constitute a formal test of a complete causal moderated-mediation process.

An additional factorial model predicted change from consensus cost, information change, centered lTPJ IBS, all three two-way interactions, and their three-way interaction. Sharedness and the sharedness × IBS interaction were subsequently added to examine whether the factorial terms and the sharedness-based model captured overlapping variation in updating. These models provided a supplementary integration of the predictors rather than a separate test of the proposed causal process.

Neural interaction estimates were evaluated using dyad-clustered CR2 covariance estimates and restricted wild-cluster-bootstrap-t tests. The sharedness × lTPJ IBS model used coefficient-specific Satterthwaite degrees of freedom. For the information-change × lTPJ IBS interaction predicting gap, CR2 inference used a t reference distribution with 20 degrees of freedom. Regional comparison models and family-wise adjustments are reported alongside the corresponding neural results.

#### 2.2.5 Results

Behavioral and IBS data were available from all 21 dyads. After blocks belonging to the same experimental cell were aggregated, the primary dataset contained 173 usable dyad × consensus-cost × information-change cells (of 189 possible cells, 16 had no available value).

Before examining updating, we tested whether the cost conditions differed in their behavioural starting points at the end of Stage 1. Categorical mixed-effects models with a dyad random intercept did not detect a significant cost effect on the perceived-consensus position, *F*(2, 144.62) = 0.13, *p* = .881; the distance between the members’ final decisions, *F*(2, 144.40) = 1.21, *p* = .301; their distance from the midpoint of the response scale, *F*(2, 155.41) = 0.53, *p* = .589; or the scale space available for movement in the direction of the new information, *F*(2, 144.38) = 2.90, *p* = .058.

##### 2.2.5.1 Behavioral results

###### Behavioral consequences of consensus-formation cost

We next examined how consensus-formation cost was associated with the three behavioral measures used in the subsequent analyses: baseline-adjusted update magnitude (*change*), perceived consensus sharedness, and between-member updating discrepancy (*gap*). These measures captured different consequences of consensus formation: how much the dyad subsequently updated, how strongly the preceding consensus was perceived to represent both members, and how similar the members’ first updates were (Figure 5).

**Figure 5.**
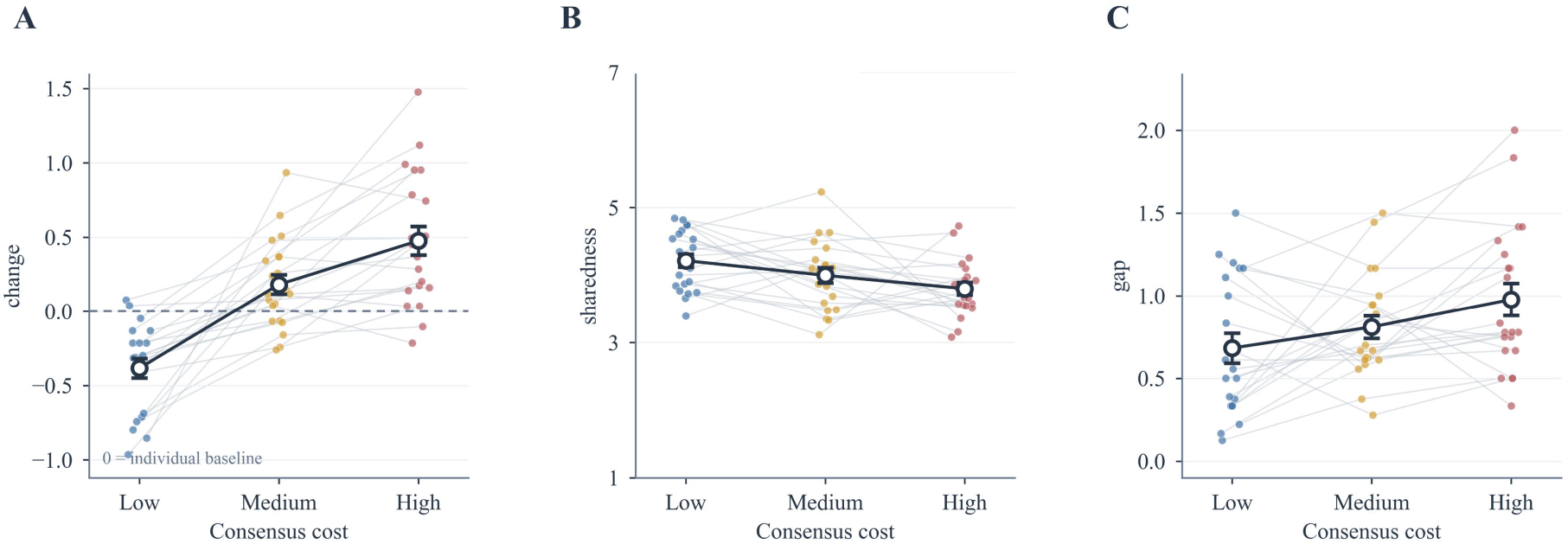
Behavioral consequences of consensus-formation cost. (A) Baseline-adjusted updating, change. The dashed line indicates equality with the information-matched individual baseline. (B) Perceived consensus sharedness. (C) Between-member updating discrepancy, gap. Small points and connecting lines show individual dyads; large points and error bars show the mean ± SE.

The same 3 × 3 categorical mixed-effects model was fitted to each outcome, with consensus cost, information change, and their interaction as fixed effects and a dyad random intercept. Consensus cost affected *change*, *F*(2, 145.16) = 53.98, *p* < .001 (Figure 5A). Using dyad-level means averaged across information conditions, the direction of the deviation from the individual baseline reversed across cost conditions. Following low-cost consensus, dyads updated less than the individual baseline (*M* = −0.383, *SD* = 0.300), *t*(20) = −5.86, *p* < .001, *d_z_* = −1.28. Following medium-cost consensus, updating was greater than the baseline (*M* = 0.182, *SD* = 0.302), *t*(20) = 2.77, *p* = .012, *d_z_* = 0.60. Following high-cost consensus, updating was further above the baseline (*M* = 0.476, *SD* = 0.443), *t*(20) = 4.93, *p* < .001, *d_z_* = 1.08. Information change also affected *change*, *F*(2, 146.19) = 31.94, *p* < .001, with smaller baseline-adjusted deviations at higher information-change levels. Because *change* subtracts the information-matched individual baseline, this effect concerns the deviation from that benchmark. The cost × information-change interaction was not significant, *F*(4, 145.59) = 0.35, *p* = .844.

Consensus cost also affected perceived sharedness, *F*(2, 147.19) = 4.92, *p* = .009 (Figure 5B). Sharedness decreased from low cost (*M* = 4.221, *SD* = 0.662) to medium cost (*M* = 3.972, *SD* = 0.763) and high cost (*M* = 3.813, *SD* = 0.693). The centered linear cost contrast was negative, *b* = −0.204, *SE* = 0.073, *p* = .005. Neither the information-change main effect nor its interaction with cost was significant (*p* = .562 and .284, respectively). Because sharedness was measured before new information appeared, these latter comparisons served as balance checks. Higher consensus-formation cost was associated with lower perceived sharedness.

The initial categorical model also identified a cost effect on *gap*, *F*(2, 145.31) = 3.58, *p* = .030 (Figure 5C). Neither the information-change main effect, *F*(2, 146.43) = 0.56, *p* = .573, nor the cost × information-change interaction, *F*(4, 145.78) = 2.17, *p* = .075, was significant. We then controlled mean absolute movement, (|*Δ_A_*| + |*Δ_B_*|)/2, to test whether the cost effect on *gap* remained after accounting for how much the two members updated overall. After this adjustment, the cost effect was no longer significant, *F*(2, 155.39) = 0.17, *p* = .841. Further adjustment for directional headroom—the scale space available in the direction of the new information—also yielded no significant cost effect, *F*(2, 154.18) = 0.05, *p* = .947. This additional control accounted for differences in the room available to update before reaching the response-scale boundary. Both covariates were retained in the subsequent neural models of *gap*.

###### Alternative explanations for the cost effect on updating

We next examined whether the association between consensus-formation cost and updating could be explained by the number of Stage 1 rounds. Round count alone predicted change, *b* = 0.124, *p* < .001. After assigned cost level was included, the round-count coefficient was close to zero, *b* = −0.003, *p* = .959, whereas the cost coefficient remained positive after controlling round count, *b* = 0.404, *p* = .013. Thus, round count did not explain additional variation beyond the assigned cost levels in this model. Because the cost categories were themselves defined by ranges of round counts, this comparison does not isolate the effect of elapsed time or repeated disagreement from formation cost.

We also tested whether the cost association changed as the experiment progressed. Neither block number, *b* = −0.006, *p* = .725, nor the cost × block-number interaction, *b* = −0.013, *p* = .092, was significant. There was therefore no clear evidence that the cost association became stronger over successive experimental blocks.

We next considered the partner’s decision adjustments during consensus formation. Mean absolute adjustment between successive Stage 1 rounds was 0.670, 0.720, and 0.794 across low-, medium-, and high-cost conditions, with no significant cost difference (*p* = .347; corresponding comparison for the other member, *p* = .333). The partner’s earlier adjustments were not significantly associated with the focal member’s subsequent update magnitude (*p* = .654) or baseline-adjusted updating (*p* = .616). These analyses did not support the size of the partner’s earlier adjustments as an explanation of the cost-related updating pattern.

Finally, an auxiliary member-level analysis tested whether the cost association was confined to the member whose update was farther from the individual baseline. Direction-aligned errors were calculated as *e_i_* = dir × *Δ_i_* − *I*⁎, and members were assigned to near and far roles separately within each block according to the absolute size of these errors. Both roles showed positive cost slopes: *e_near_*, *b* = 0.227, *SE* = 0.046, *p* < .001; *e_far_*, *b* = 0.584, *SE* = 0.101, *p* < .001. Randomly assigning the two members to the roles 1,000 times produced an unsorted reference centred at 0.406, with a 95% reference interval of [0.325, 0.486]. The near and far slopes fell below and above this interval, respectively, consistent with the sorting procedure separating their estimates. Both slopes remained positive, indicating that the cost association was present in both member roles.

###### Perceived sharedness linked consensus cost to updating

Consensus cost was negatively associated with perceived sharedness, *a* = −0.204, CR2 *SE* = 0.072, *p* = .011. Controlling for cost and information change, higher sharedness predicted less updating, *b* = −0.195, CR2 *SE* = 0.059, *p* = .005. The indirect association was *ab* = 0.040, joint-significance *p* = .011, corresponding to 9.3% of the total association. The total cost coefficient was *c* = 0.430, CR2 *SE* = 0.052, *p* < .001; after sharedness was included, the direct coefficient remained positive, *c′* = 0.390, CR2 *SE* = 0.048, *p* < .001 (Figure 6). The dyad-bootstrap 95% confidence interval for ab was [0.009, 0.075].

**Figure 6.**
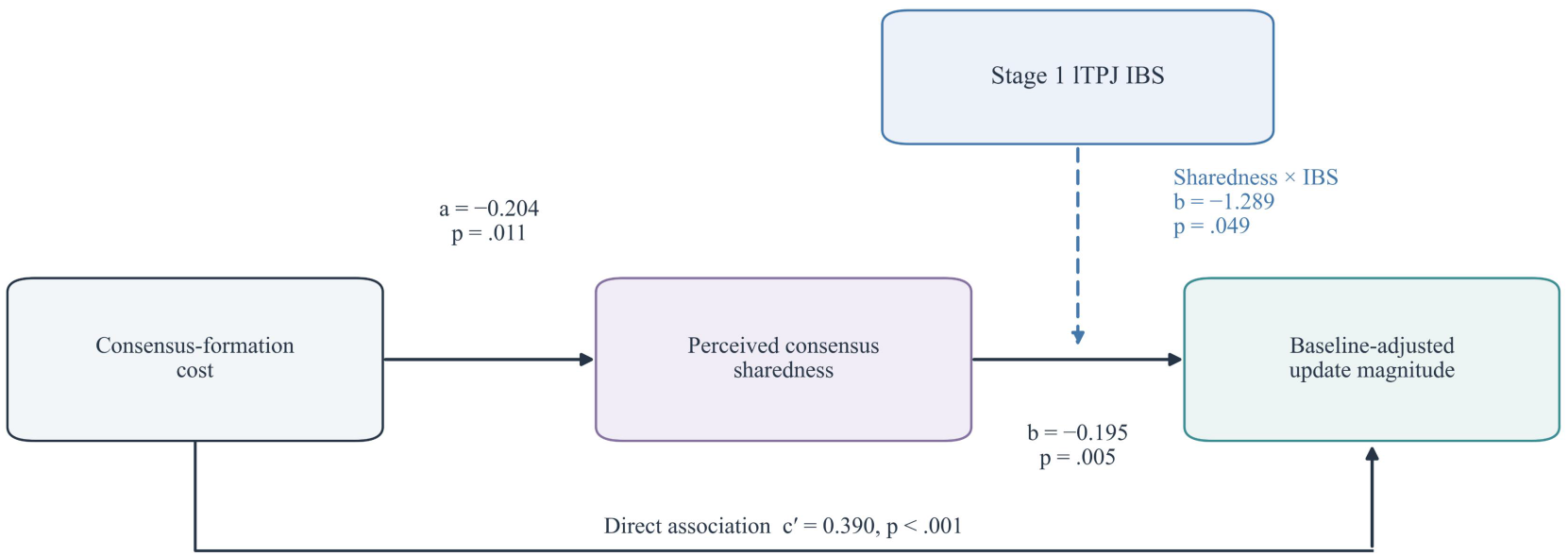
Sharedness pathway and neural moderation of updating.

##### 2.2.5.2 Neural results

###### Frequency selection and properties of Stage 1 IBS

The whole-channel Stage 1 frequency scan identified two significant clusters for the effect of consensus-formation cost (Figure 7). The higher-frequency cluster, approximately 0.16–0.18 Hz, overlapped a systemic physiological-frequency range and was not carried forward. The retained cluster spanned 0.019–0.023 Hz. Its observed cluster F-sum was 17.424; 2 of the 1,000 permuted maximum cluster F-sums equaled or exceeded this value, giving a cluster-level *p* = .002. Figure 7 shows the channel-by-frequency F map and the frequency-wise whole-channel statistics.

**Figure 7.**
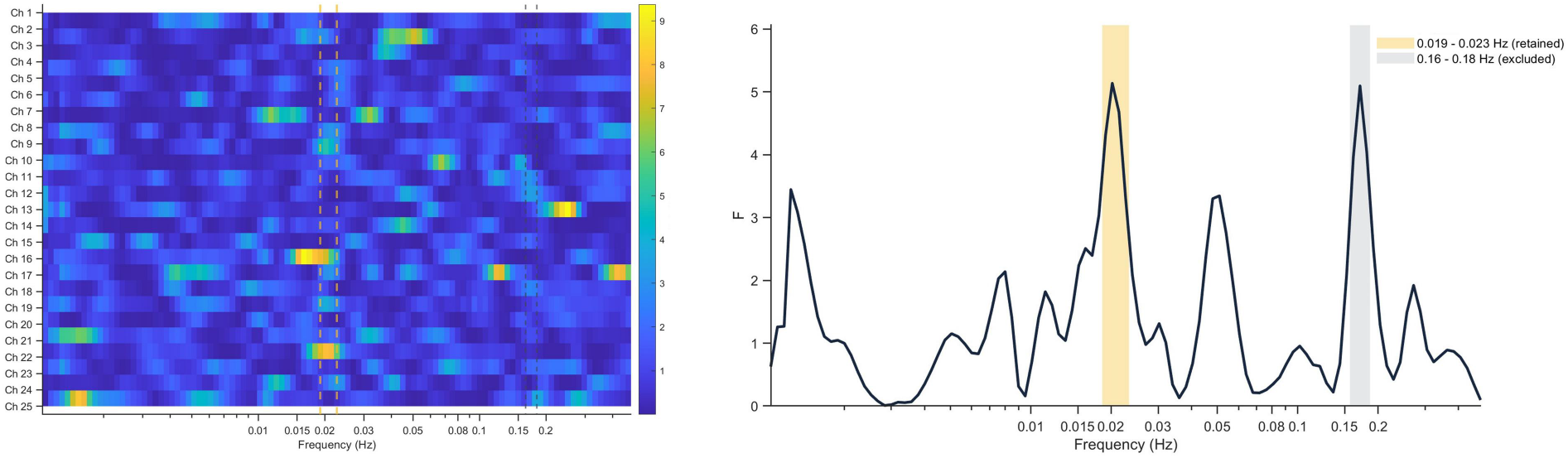
Frequency selection for inter-brain synchrony during consensus formation. Inter-brain synchrony (IBS) was estimated from oxygenated haemoglobin signals using wavelet transform coherence during Stage 1 in 21 dyads. Left, channel-by-frequency F statistics for the effect of consensus-formation cost; colour indicates F values, and dashed lines delimit the highlighted frequency ranges. Right, frequency-wise F statistics from repeated-measures analyses of variance on whole-channel mean Fisher-z-transformed coherence, with consensus cost as the within-dyad factor. Yellow shading identifies the retained 0.019–0.023 Hz cluster (cluster-level *p* = .002). Grey shading identifies the 0.16–0.18 Hz cluster, which was excluded because of its overlap with a systemic physiological frequency range.

The selected band was applied to all regional analyses. Within this band, whole-channel Fisher-z IBS means were 0.671, 0.637, and 0.638 across low, medium, and high cost, respectively, *F*(2, 40) = 4.71, *p* = .015. Because frequency selection was based on the whole-channel cost contrast, this result describes the selected band rather than providing an independent test of the contrast.

For lTPJ, IBS means were 0.670, 0.658, and 0.657, with no significant cost effect, *F*(2, 40) = 0.25, *p* = .776. All subsequent neural–behavioral models used Fisher-z IBS values centered separately within each consensus-cost condition. These lTPJ IBS values were not significantly associated with sharedness (*r* = .109; dyad-clustered CR2/Satterthwaite test of the regression slope, *p* = .078).

###### lTPJ IBS was associated with smaller updating discrepancies under large information changes

We first examined whether Stage 1 lTPJ IBS was associated with the similarity of the members’ subsequent updates. The information-change × lTPJ IBS interaction predicting gap was negative, *b* = −1.684, 95% CI [−3.071, −0.297], CR2 *p* = .0198; restricted wild-cluster-bootstrap *p* = .0076. The fitted association between lTPJ IBS and gap became more negative as information change increased: under larger information changes, higher lTPJ IBS was associated with smaller discrepancies between the members’ first updates. The six-measure regional comparison is reported in **Table S2**.

###### Sharedness and lTPJ IBS jointly predicted updating

We then tested whether lTPJ IBS moderated the association between perceived sharedness and update magnitude (Figure 6). The sharedness × lTPJ IBS interaction predicting change was negative, *b* = −1.289, CR2 *SE* = 0.426, *t*(3.36) = −3.03, *p* = .049, 95% CI [−2.566, −0.013], using Satterthwaite degrees of freedom. The restricted wild-cluster-bootstrap test yielded *p* = .0578, indicating sensitivity to the inference method.

The estimated sharedness slope became more negative as lTPJ IBS increased (Figure 8). At −1 SD IBS, the slope was −0.020, 95% CI [−0.192, 0.151]; at mean IBS, −0.145, 95% CI [−0.263, −0.026]; and at +1 SD IBS, −0.269, 95% CI [−0.418, −0.120]. Thus, higher sharedness was associated with less updating at mean and high lTPJ IBS, whereas this association was not detected at low IBS. At mean sharedness, the conditional IBS slope was not significant, *b* = −0.362, 95% CI [−1.797, 1.073], *p* = .565. The corresponding regional comparisons are reported in **Table S2**.

**Figure 8.**
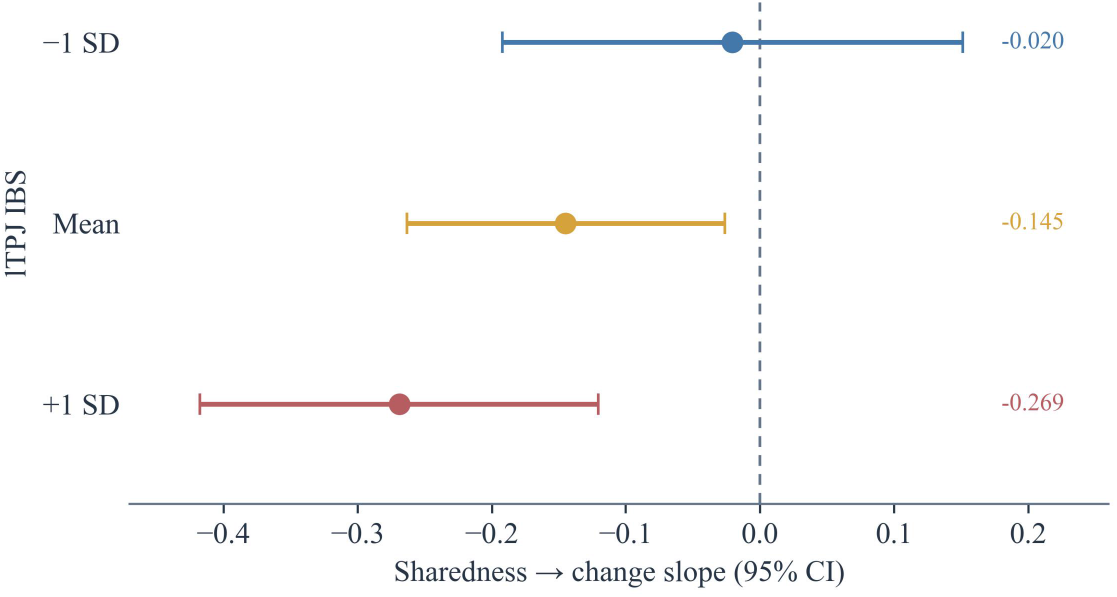
Conditional associations between perceived sharedness and baseline-adjusted updating. Horizontal error bars show 95% CR2/Satterthwaite confidence intervals.

## 3. Discussion

The history of reaching agreement shaped how dyads responded when information changed. After low-cost consensus, dyads revised less than individuals working alone; after medium- and high-cost consensus, they revised more. Greater formation cost reduced the extent to which members felt that the established judgment represented both of them, and lower perceived sharedness was associated with greater updating. This indirect association accounted for a modest part of the cost effect and the estimated sharedness–updating association was stronger at higher lTPJ IBS. Higher IBS also predicted more similar signed updatings when information changed substantially. Together, these findings support an account in which costly consensus formation loosens an agreement’s hold partly by weakening perceived joint endorsement, while interpersonal alignment shapes both the influence of that endorsement and the coordination of subsequent updating.

### 3.1 Formation history and the psychological status of consensus

The crossover relative to individual updating suggests that consensus did not exert a fixed influence on subsequent judgment. Dyads updated less than individuals after a low-cost consensus, but more after medium- and high-cost consensus. This pattern is difficult to reconcile with a simple investment-based account, under which prior expenditure strengthens persistence with an established choice (Arkes & Blumer, 1985; Whyte, 1993). Instead, formation history appears to change the psychological status of the agreement itself.

A useful distinction is between the effort invested in reaching agreement and the social information acquired along the way. The former concerns what members have spent; the latter concerns what they have learned about the support for their judgment. These consequences need not strengthen commitment in the same direction. Agreement can increase confidence, whereas disagreement can reduce it (Pescetelli & Yeung, 2020; Kappes et al., 2020). Similarly, groups with more homogeneous initial preferences show a stronger preference for supporting information, an effect mediated by greater confidence and commitment (Schulz-Hardt et al., 2000). Formation history may therefore influence later updating by changing the basis on which the existing judgment is retained, rather than merely adding a cost to abandoning it.

At low formation cost, the established judgment may acquire both epistemic and relational value. Shared-reality theory links perceived commonality to subjective certainty and social connection: a judgment experienced as shared provides reassurance about the world while affirming a connection with another person (Higgins et al., 2021; Echterhoff et al., 2009; Rossignac-Milon et al., 2021). Revising it may therefore entail more than changing an individual estimate; it may put a successfully established common understanding at risk. This provides a motivational basis for preserving an easily reached consensus. It also suggests a route through information processing. Confidence can selectively reduce the integration of evidence inconsistent with an existing choice (Rollwage et al., 2020). To the extent that easy agreement provides such reassurance, members may give the established judgment greater weight when circumstances change. Its persistence would then reflect the value of the shared state being maintained, not the effort required to achieve it.

High-cost consensus requires the converse process to be specified more carefully. Merely removing the additional reassurance of agreement would predict a return toward individual updating, not necessarily updating beyond it. Repeated failures to agree may instead introduce doubt that would not arise when judging alone. Even after agreement is announced, the preceding difficulty may remain evidence that the settled position is not securely supported by a common understanding. On this account, costly formation does not simply fail to strengthen the judgment; it can weaken members’ confidence in using it as the continuing basis for choice. New information would then encounter an already unsettled judgment and receive greater relative weight. The finding that disagreement can reduce confidence below its pre-interaction level provides a precedent for this possibility, although its persistence after an eventual consensus remains a hypothesis of the present account (Pescetelli & Yeung, 2020).

This interpretation places consensus updating within a temporally extended process of social validation. The formation phase establishes not only a response but also how securely that response is experienced as jointly supported; the subsequent information change tests a judgment whose standing has already been altered by that history. Our sharedness findings are consistent with this distinction, without establishing that sharedness and confidence are interchangeable. A discriminating prediction follows: equally lengthy delays attributed to a nonsocial obstacle should not produce the same updating pattern as repeated indications of disagreement. Conversely, credible confirmation of both members’ independent endorsement should attenuate the effect of a difficult formation history even when the effort already expended remains unchanged. The theoretical implication is that agreement can end negotiation without erasing the social information generated during it. That information may continue to determine whether the resulting judgment is protected or reopened when the world changes.

### 3.2 Interpersonal alignment in consensus maintenance and updating

An agreement can be experienced as jointly endorsed without specifying how its members will respond when circumstances change. Two people may support the same judgment yet differ in what they regard as sufficient grounds for revising it. Joint-action accounts emphasize that coordinated behavior depends not only on shared representations but also on anticipating a partner’s actions and integrating them with one’s own (Sebanz et al., 2006). Applied to consensus updating, this suggests that members face two related questions: how the new information changes their judgment, and how their response will fit with the partner’s. We propose that interpersonal alignment helps connect the subjective status of an established agreement to expectations about its subsequent use.

This distinction provides a process account of the stronger estimated association between sharedness and reduced updating at higher lTPJ IBS, although the statistical support for this interaction was inference-sensitive. When a judgment feels representative of both members, each may expect the other to continue giving it substantial weight. Retaining it then preserves not only a position considered valid but also a response expected to remain compatible with the partner’s. Greater alignment may make perceived joint endorsement a more informative guide to the partner’s likely response, strengthening this additional reason for restraint. With weaker alignment, knowing that a judgment was previously shared may provide less guidance about how the partner will respond to changed circumstances. On this account, alignment does not simply increase commitment; it strengthens the connection between perceived endorsement and anticipated behavior.

The updating-discrepancy result helps distinguish this account from simple inertia. Under larger information changes, higher formation-period lTPJ IBS was associated with more similar signed updates, in a model controlling for mean movement and directional headroom. This association therefore concerns the compatibility of members’ adjustments, rather than merely their reluctance to move. Predictive accounts of social alignment emphasize detecting and reducing interpersonal mismatch (Shamay-Tsoory et al., 2019). Extending this logic, preserving compatibility after substantial change may require departing from the established judgment together rather than defending it. Alignment may support comparable weighting of the changed evidence and expectations of similar adjustments. The relevant commonality is then not an unchanged answer, but a compatible way of transforming the previous answer. Indeed, members in the present task could make matching signed updates from different perceived consensus values.

The timing of the association further suggests that coordination may operate prospectively rather than only through reactions to observed disagreement. IBS was measured during formation, whereas the first updates were submitted before renewed agreement feedback. Their correspondence therefore cannot be attributed to convergence produced by feedback on those updates. Accounts of brain-to-brain coupling propose that interaction constrains how individuals organize joint behavior (Hasson et al., 2012).

Experimental evidence also shows that social exchanges of confidence can alter subjective uncertainty and neural signatures of evidence accumulation (Esmaily et al., 2023), indicating that social influence can enter the decision process rather than merely modify its reported outcome. The present findings motivate a temporal extension: coordination established during one decision may remain relevant to how members approach the next, even after the evidence supporting their earlier agreement has changed.

Anticipation of the partner’s response remains a proposed mechanism, not a computation directly identified by IBS (Hamilton, 2021; Holroyd, 2022). Compatible processing of shared task information could also contribute to the observed associations. A discriminating test would separate private updating from updating under a renewed requirement to agree. Following identical formation histories, the social validation of the original judgment might continue to influence private responses, whereas an additional influence arising from anticipated coordination should weaken when members no longer need to match their adjustments. Eliciting expectations about the partner’s update would further test whether these expectations account for the difference between private and joint responses. This would distinguish retaining a socially validated judgment from retaining it partly because the partner is expected to do so too.

Formation history may thus leave two distinguishable legacies: an evaluation of how securely a judgment is jointly supported, and expectations about how to respond as a pair. The former concerns the standing of the established judgment; the latter concerns the organization of subsequent behavior. Their interaction offers an explanation for how consensus can support both persistence and change: maintaining coordination need not require maintaining the content of the agreement.

## Data Availability

The data supporting the findings of this study are available from the corresponding author upon reasonable request.

## References

Arkes, H. R., & Blumer, C. (1985). The psychology of sunk cost. Organizational Behavior and Human Decision Processes, 35(1), 124–140. 10.1016/0749-5978(85)90049-4

Badre, D., & D’Esposito, M. (2009). Is the rostro-caudal axis of the frontal lobe hierarchical? Nature Reviews Neuroscience, 10(9), 659–669. 10.1038/nrn2667

Bahrami, B., Olsen, K., Latham, P. E., Roepstorff, A., Rees, G., & Frith, C. D. (2010). Optimally interacting minds. Science, 329(5995), 1081–1085. 10.1126/science.1185718

Bell, R. M., & McCaffrey, D. F. (2002). Bias reduction in standard errors for linear regression with multi-stage samples. Survey Methodology, 28(2), 169–181.

Conradt, L., & Roper, T. J. (2005). Consensus decision making in animals. Trends in Ecology & Evolution, 20(8), 449–456. 10.1016/j.tree.2005.05.008

Cope, M., & Delpy, D. T. (1988). System for long-term measurement of cerebral blood and tissue oxygenation on newborn infants by near infra-red transillumination. Medical & Biological Engineering & Computing, 26(3), 289–294. 10.1007/BF02447083

Cui, X., Bryant, D. M., & Reiss, A. L. (2012). NIRS-based hyperscanning reveals increased interpersonal coherence in superior frontal cortex during cooperation. NeuroImage, 59(3), 2430–2437. 10.1016/j.neuroimage.2011.09.003

Czeszumski, A., Liang, S. H.-Y., Dikker, S., König, P., Lee, C.-P., Koole, S. L., & Kelsen, B. (2022). Cooperative behavior evokes interbrain synchrony in the prefrontal and temporoparietal cortex: A systematic review and meta-analysis of fNIRS hyperscanning studies. eNeuro, 9(2), Article ENEURO.0268-21.2022. 10.1523/ENEURO.0268-21.2022

Deng, W., Garg, K., & Mobbs, D. (2026). Linking compromise and responsibility attribution to risky decision-making in dyadic foraging. Proceedings of the National Academy of Sciences, 123(27), Article e2602570123. 10.1073/pnas.2602570123

Echterhoff, G., & Higgins, E. T. (2017). Creating shared reality in interpersonal and intergroup communication: The role of epistemic processes and their interplay. European Review of Social Psychology, 28(1), 175–226. 10.1080/10463283.2017.1333315

Echterhoff, G., Higgins, E. T., & Levine, J. M. (2009). Shared reality: Experiencing commonality with others’ inner states about the world. Perspectives on Psychological Science, 4(5), 496–521. 10.1111/j.1745-6924.2009.01161.x

Esmaily, J., Zabbah, S., Ebrahimpour, R., & Bahrami, B. (2023). Interpersonal alignment of neural evidence accumulation to social exchange of confidence. eLife, 12, Article e83722. 10.7554/eLife.83722

Grinsted, A., Moore, J. C., & Jevrejeva, S. (2004). Application of the cross wavelet transform and wavelet coherence to geophysical time series. Nonlinear Processes in Geophysics, 11(5/6), 561–566. 10.5194/npg-11-561-2004

Hamilton, A. F. de C. (2021). Hyperscanning: Beyond the hype. Neuron, 109(3), 404–407. 10.1016/j.neuron.2020.11.008

Hasson, U., Ghazanfar, A. A., Galantucci, B., Garrod, S., & Keysers, C. (2012). Brain-to-brain coupling: A mechanism for creating and sharing a social world. Trends in Cognitive Sciences, 16(2), 114–121. 10.1016/j.tics.2011.12.007

Higgins, E. T., Rossignac-Milon, M., & Echterhoff, G. (2021). Shared reality: From sharing-is-believing to merging minds. Current Directions in Psychological Science, 30(2), 103–110. 10.1177/0963721421992027

Holroyd, C. B. (2022). Interbrain synchrony: On wavy ground. Trends in Neurosciences, 45(5), 346–357. 10.1016/j.tins.2022.02.002

Jiang, J., Chen, C., Dai, B., Shi, G., Ding, G., Liu, L., & Lu, C. (2015). Leader emergence through interpersonal neural synchronization. Proceedings of the National Academy of Sciences, 112(14), 4274–4279. 10.1073/pnas.1422930112

Jurcak, V., Tsuzuki, D., & Dan, I. (2007). 10/20, 10/10, and 10/5 systems revisited: Their validity as relative head-surface-based positioning systems. NeuroImage, 34(4), 1600–1611. 10.1016/j.neuroimage.2006.09.024

Kappes, A., Harvey, A. H., Lohrenz, T., Montague, P. R., & Sharot, T. (2020). Confirmation bias in the utilization of others’ opinion strength. Nature Neuroscience, 23(1), 130–137. 10.1038/s41593-019-0549-2

Kerr, N. L., & Tindale, R. S. (2004). Group performance and decision making. Annual Review of Psychology, 55, 623–655. 10.1146/annurev.psych.55.090902.142009

Krause, J., & Ruxton, G. D. (2002). Living in groups. Oxford University Press.

Kriegeskorte, N., Simmons, W. K., Bellgowan, P. S. F., & Baker, C. I. (2009). Circular analysis in systems neuroscience: The dangers of double dipping. Nature Neuroscience, 12(5), 535–540. 10.1038/nn.2303

MacKinnon, D. P., Lockwood, C. M., Hoffman, J. M., West, S. G., & Sheets, V. (2002). A comparison of methods to test mediation and other intervening variable effects. Psychological Methods, 7(1), 83–104. 10.1037/1082-989X.7.1.83

Montague, P. R., Berns, G. S., Cohen, J. D., McClure, S. M., Pagnoni, G., Dhamala, M., Wiest, M. C., Karpov, I., King, R. D., Apple, N., & Fisher, R. E. (2002). Hyperscanning: Simultaneous fMRI during linked social interactions. NeuroImage, 16(4), 1159–1164. 10.1006/nimg.2002.1150

Pescetelli, N., & Yeung, N. (2020). The effects of recursive communication dynamics on belief updating. Proceedings of the Royal Society B: Biological Sciences, 287(1931), Article 20200025. 10.1098/rspb.2020.0025

Pustejovsky, J. E., & Tipton, E. (2018). Small-sample methods for cluster-robust variance estimation and hypothesis testing in fixed effects models. Journal of Business & Economic Statistics, 36(4), 672– 683. 10.1080/07350015.2016.1247004

Redcay, E., & Schilbach, L. (2019). Using second-person neuroscience to elucidate the mechanisms of social interaction. Nature Reviews Neuroscience, 20(8), 495–505. 10.1038/s41583-019-0179-4

Reinero, D. A., Dikker, S., & Van Bavel, J. J. (2021). Inter-brain synchrony in teams predicts collective performance. Social Cognitive and Affective Neuroscience, 16(1–2), 43–57. 10.1093/scan/nsaa135

Rollwage, M., Loosen, A., Hauser, T. U., Moran, R., Dolan, R. J., & Fleming, S. M. (2020). Confidence drives a neural confirmation bias. Nature Communications, 11(1), Article 2634. 10.1038/s41467-020-16278-6

Rossignac-Milon, M., Bolger, N., Zee, K. S., Boothby, E. J., & Higgins, E. T. (2021). Merged minds: Generalized shared reality in dyadic relationships. Journal of Personality and Social Psychology, 120(4), 882–911. 10.1037/pspi0000266

Samson, D., Apperly, I. A., Chiavarino, C., & Humphreys, G. W. (2004). Left temporoparietal junction is necessary for representing someone else’s belief. Nature Neuroscience, 7(5), 499–500. 10.1038/nn1223

Saxe, R., & Kanwisher, N. (2003). People thinking about thinking people: The role of the temporo-parietal junction in "theory of mind." NeuroImage, 19(4), 1835–1842. 10.1016/S1053-8119(03)00230-1

Schulz-Hardt, S., Frey, D., Lüthgens, C., & Moscovici, S. (2000). Biased information search in group decision making. Journal of Personality and Social Psychology, 78(4), 655–669. 10.1037/0022-3514.78.4.655

Schurz, M., Radua, J., Aichhorn, M., Richlan, F., & Perner, J. (2014). Fractionating theory of mind: A meta-analysis of functional brain imaging studies. Neuroscience & Biobehavioral Reviews, 42, 9–34. 10.1016/j.neubiorev.2014.01.009

Sebanz, N., Bekkering, H., & Knoblich, G. (2006). Joint action: Bodies and minds moving together. Trends in Cognitive Sciences, 10(2), 70–76. 10.1016/j.tics.2005.12.009

Shamay-Tsoory, S. G., Saporta, N., Marton-Alper, I. Z., & Gvirts, H. Z. (2019). Herding brains: A core neural mechanism for social alignment. Trends in Cognitive Sciences, 23(3), 174–186. 10.1016/j.tics.2019.01.002

Singh, A. K., Okamoto, M., Dan, H., Jurcak, V., & Dan, I. (2005). Spatial registration of multichannel multi-subject fNIRS data to MNI space without MRI. NeuroImage, 27(4), 842–851. 10.1016/j.neuroimage.2005.05.019

Sleesman, D. J., Conlon, D. E., McNamara, G., & Miles, J. E. (2012). Cleaning up the big muddy: A meta-analytic review of the determinants of escalation of commitment. Academy of Management Journal, 55(3), 541–562. 10.5465/amj.2010.0696

Spencer, S. J., Zanna, M. P., & Fong, G. T. (2005). Establishing a causal chain: Why experiments are often more effective than mediational analyses in examining psychological processes. Journal of Personality and Social Psychology, 89(6), 845–851. 10.1037/0022-3514.89.6.845

Staw, B. M. (1976). Knee-deep in the big muddy: A study of escalating commitment to a chosen course of action. Organizational Behavior and Human Performance, 16(1), 27–44. 10.1016/0030-5073(76)90005-2

Suzuki, S., Adachi, R., Dunne, S., Bossaerts, P., & O’Doherty, J. P. (2015). Neural mechanisms underlying human consensus decision-making. Neuron, 86(2), 591–602. 10.1016/j.neuron.2015.03.019

Tachtsidis, I., & Scholkmann, F. (2016). False positives and false negatives in functional near-infrared spectroscopy: Issues, challenges, and the way forward. Neurophotonics, 3(3), Article 031405. 10.1117/1.NPh.3.3.031405

Tang, H., Mai, X., Wang, S., Zhu, C., Krueger, F., & Liu, C. (2016). Interpersonal brain synchronization in the right temporo-parietal junction during face-to-face economic exchange. Social Cognitive and Affective Neuroscience, 11(1), 23–32. 10.1093/scan/nsv092

Tsuzuki, D., Jurcak, V., Singh, A. K., Okamoto, M., Watanabe, E., & Dan, I. (2007). Virtual spatial registration of stand-alone fNIRS data to MNI space. NeuroImage, 34(4), 1506–1518. 10.1016/j.neuroimage.2006.10.043

Whyte, G. (1993). Escalating commitment in individual and group decision making: A prospect theory approach. Organizational Behavior and Human Decision Processes, 54(3), 430–455. 10.1006/obhd.1993.1018

Yzerbyt, V., Muller, D., Batailler, C., & Judd, C. M. (2018). New recommendations for testing indirect effects in mediational models: The need to report and test component paths. Journal of Personality and Social Psychology, 115(6), 929–943. 10.1037/pspa0000132

