## Supplementary material for "Consensus Formation History Shapes Collective Updating": Table S2

**Regional comparison of inter-brain synchrony interactions in post-consensus updating**

Information change × IBS predicting between-member updating discrepancy (gap)

| **ROI** | **b** | **CR2 p** | **WCB p** | **Bonferroni p (WCB)** |
| --- | --- | --- | --- | --- |
| lTPJ | -1.684 | .0198 | .0076 | .0456 |
| rTPJ | -0.013 | .992 | .993 | 1.000 |
| aDPFC | -1.388 | .218 | .253 | 1.000 |
| mDPFC | -1.190 | .295 | .315 | 1.000 |
| pDPFC | -0.583 | .678 | .686 | 1.000 |
| Whole-brain | -2.308 | .198 | .230 | 1.000 |

Perceived consensus sharedness × IBS predicting baseline-adjusted updating (change)

| **ROI** | **b** | **CR2 p** | **WCB p** | **Bonferroni p (WCB)** |
| --- | --- | --- | --- | --- |
| lTPJ | -1.289 | .049 | .0578 | .3468 |
| rTPJ | -0.270 | .809 | .798 | 1.000 |
| aDPFC | -1.776 | .0176 | .0453 | .2718 |
| mDPFC | 0.058 | .935 | .928 | 1.000 |
| pDPFC | -0.411 | .636 | .627 | 1.000 |
| Whole-brain | -1.627 | .125 | .116 | .696 |

*Note.* Coefficients are unstandardized. IBS values were centered within each consensus-cost condition. Panel A models controlled mean absolute movement and directional headroom. Panel B models included consensus-formation cost, information change, perceived consensus sharedness, IBS, and the sharedness × IBS interaction. CR2 p values use dyad-clustered CR2 covariance estimates with Satterthwaite-adjusted degrees of freedom; WCB p values use restricted wild-cluster-bootstrap-t tests. Bonferroni was calculated separately within each panel across five anatomical ROIs and the whole-brain summary. lTPJ was the literature-informed primary ROI; the adjusted p values evaluate regional specificity.
