## Supplementary material for "Consensus Formation History Shapes Collective Updating": Table S1

**Supplementary Material 1**

Locations on the International 10/05 system, as well as the corresponding anatomical labels, region of interest and MNI coordinates for the placement of fNIRS channels.

| **Channel** | **Position** | **X** | **Y** | **Z** | **Anatomical Label** |
| --- | --- | --- | --- | --- | --- |
| 1 | Fp1h | -12.07507 | 71.036253 | -4.843375 | anterior dorsal prefrontal cortex |
| 2 | Fp2h | 14.23633 | 70.823186 | -5.301705 | anterior dorsal prefrontal cortex |
| 3 | AFp5 | -27.64447 | 66.512257 | -1.937112 | anterior dorsal prefrontal cortex |
| 4 | AFpz | 2.583211 | 68.870082 | 10.934657 | anterior dorsal prefrontal cortex |
| 5 | AFp6 | 30.390812 | 66.99523 | -1.662572 | anterior dorsal prefrontal cortex |
| 6 | AF1 | -14.70924 | 65.626723 | 22.912322 | anterior dorsal prefrontal cortex |
| 7 | AF2 | 16.915452 | 66.827437 | 23.990586 | anterior dorsal prefrontal cortex |
| 8 | AFF3h | -26.01439 | 55.55286 | 30.528652 | mid-dorsal prefrontal cortex |
| 9 | AFFz | 1.574451 | 55.723029 | 38.394758 | mid-dorsal prefrontal cortex |
| 10 | AFF4h | 28.786208 | 55.927454 | 31.030869 | mid-dorsal prefrontal cortex |
| 11 | F1h | -10.49759 | 45.338513 | 50.740331 | mid-dorsal prefrontal cortex |
| 12 | F2h | 12.859702 | 45.450615 | 50.666019 | mid-dorsal prefrontal cortex |
| 13 | FFC1 | -23.29506 | 31.671745 | 54.929585 | posterior dorsal prefrontal cortex |
| 14 | FFCz | 2.255396 | 30.64247 | 57.643211 | posterior dorsal prefrontal cortex |
| 15 | FFC2 | 25.903854 | 31.404311 | 55.886701 | posterior dorsal prefrontal cortex |
| 16 | FC1h | -13.6313 | 17.580136 | 67.405904 | posterior dorsal prefrontal cortex |
| 17 | FC2h | 14.886146 | 17.230097 | 66.576688 | posterior dorsal prefrontal cortex |
| 18 | CP5h | -61.05199 | -43.48402 | 44.456271 | left temporoparietal junction |
| 19 | CP6h | 62.090888 | -43.18651 | 44.594387 | right temporoparietal junction |
| 20 | CPP5 | -62.31091 | -56.60245 | 28.632515 | left temporoparietal junction |
| 21 | CPP3 | -49.52764 | -56.58848 | 52.636462 | left temporoparietal junction |
| 22 | CPP4 | 50.15994 | -56.479998 | 53.49844 | right temporoparietal junction |
| 23 | CPP6 | 61.838154 | -56.24619 | 28.82676 | right temporoparietal junction |
| 24 | P5h | -51.65355 | -69.21793 | 39.956249 | left temporoparietal junction |
| 25 | P6h | 50.856757 | -68.19701 | 39.647664 | right temporoparietal junction |
